# StandardRat: A multi-center consensus protocol to enhance functional connectivity specificity in the rat brain

**DOI:** 10.1101/2022.04.27.489658

**Authors:** Joanes Grandjean, Gabriel Desrosiers-Gregoire, Cynthia Anckaerts, Diego Angeles-Valdez, Fadi Ayad, David A Barrière, Ines Blockx, Aleksandra B Bortel, Margaret Broadwater, Beatriz M Cardoso, Marina Célestine, Jorge E Chavez-Negrete, Sangcheon Choi, Emma Christiaen, Perrin Clavijo, Luis Colon-Perez, Samuel Cramer, Tolomeo Daniele, Elaine Dempsey, Yujian Diao, Arno Doelemeyer, David Dopfel, Lenka Dvořáková, Claudia Falfán-Melgoza, Francisca F Fernandes, Caitlin F Fowler, Antonio Fuentes-Ibañez, Clément Garin, Eveline Gelderman, Carla EM Golden, Chao CG Guo, Marloes JAG Henckens, Lauren A Hennessy, Peter Herman, Nita Hofwijks, Corey Horien, Tudor M Ionescu, Jolyon Jones, Johannes Kaesser, Eugene Kim, Henriette Lambers, Alberto Lazari, Sung-Ho Lee, Amanda Lillywhite, Yikang Liu, Yanyan Y Liu, Alejandra López-Castro, Xavier López-Gil, Zilu Ma, Eilidh MacNicol, Dan Madularu, Francesca Mandino, Sabina Marciano, Matthew J McAuslan, Patrick McCunn, Alison McIntosh, Xianzong Meng, Lisa Meyer-Baese, Stephan Missault, Federico Moro, Daphne Naessens, Laura J Nava-Gomez, Hiroi Nonaka, Juan J Ortiz, Jaakko Paasonen, Lore M Peeters, Mickaël Pereira, Pablo D Perez, Marjory Pompilus, Malcolm Prior, Rustam Rakhmatullin, Henning M Reimann, Jonathan Reinwald, Rodrigo Triana de Rio, Alejandro Rivera-Olvera, Daniel Ruiz-Pérez, Gabriele Russo, Tobias J Rutten, Rie Ryoke, Markus Sack, Piergiorgio Salvan, Basavaraju G Sanganahalli, Aileen Schroeter, Bhedita J Seewoo, Erwan Selingue, Aline Seuwen, Bowen Shi, Nikoloz Sirmpilatze, Joanna AB Smith, Corrie Smith, Filip Sobczak, Petteri J Stenroos, Milou Straathof, Sandra Strobelt, Akira Sumiyoshi, Kengo Takahashi, Maria E Torres-García, Raul Tudela, Monica van den Berg, Kajo van der Marel, Aran TB van Hout, Roberta Vertullo, Benjamin Vidal, Roel M Vrooman, Victora X Wang, Isabel Wank, David JG Watson, Ting Yin, Yongzhi Zhang, Stefan Zurbruegg, Sophie Achard, Sarael Alcauter, Dorothee P Auer, Emmanuel L Barbier, Jürgen Baudewig, Christian F Beckmann, Nicolau Beckmann, Guillaume JPC Becq, Erwin LA Blezer, Radu Bolbos, Susann Boretius, Sandrine Bouvard, Eike Budinger, Joseph D Buxbaum, Diana Cash, Victoria Chapman, Kai-Hsiang Chuang, Luisa Ciobanu, Bram Coolen, Jeffrey W Dalley, Marc Dhenain, Rick M Dijkhuizen, Oscar Esteban, Cornelius Faber, Marcelo Febo, Kirk W Feindel, Gianluigi Forloni, Jérémie Fouquet, Eduardo A Garza-Villarreal, Natalia Gass, Jeffrey C Glennon, Alessandro Gozzi, Olli Gröhn, Andrew Harkin, Arend Heerschap, Xavier Helluy, Kristina Herfert, Arnd Heuser, Judith R Homberg, Danielle J Houwing, Fahmeed Hyder, Giovanna Diletta Ielacqua, Ileana O Jelescu, Heidi Johansen-Berg, Gen Kaneko, Ryuta Kawashima, Shella D Keilholz, Georgios A Keliris, Clare Kelly, Christian Kerskens, Jibran Y Khokhar, Peter C Kind, Jean-Baptiste Langlois, Jason P Lerch, Monica A López-Hidalgo, Denise Manahan-Vaughan, Fabien Marchand, Rogier B Mars, Gerardo Marsella, Edoardo Micotti, Emma Muñoz-Moreno, Jamie Near, Thoralf Niendorf, Willem M Otte, Patricia Pais, Wen-Ju Pan, Roberto A Prado-Alcalá, Gina L Quirarte, Jennifer Rodger, Tim Rosenow, Cassandra Sampaio Baptista, Alexander Sartorius, Stephen J Sawiak, Tom WJ Scheenen, Noam Shemesh, Yen-Yu Ian Shih, Amir Shmuel, Guadalupe Soria, Ron Stoop, Garth J Thompson, Sally M Till, Nick Todd, Annemie Van Der Linden, Annette van der Toorn, Geralda AF van Tilborg, Christian Vanhove, Andor Veltien, Marleen Verhoye, Lydia Wachsmuth, Wolfgang Weber-Fahr, Patricia Wenk, Xin Yu, Valerio Zerbi, Nanyin Zhang, Baogui B Zhang, Luc Zimmer, Gabriel A Devenyi, M Mallar Chakravarty, Andreas Hess

## Abstract

Task-free functional connectivity in animal models provides an experimental framework to examine connectivity phenomena under controlled conditions and allows comparison with invasive or terminal procedures. To date, animal acquisitions are performed with varying protocols and analyses that hamper result comparison and integration. We introduce *StandardRat*, a consensus rat functional MRI acquisition protocol tested across 20 centers. To develop this protocol with optimized acquisition and processing parameters, we initially aggregated 65 functional imaging datasets acquired in rats from 46 centers. We developed a reproducible pipeline for the analysis of rat data acquired with diverse protocols and determined experimental and processing parameters associated with a more robust functional connectivity detection. We show that the standardized protocol enhances biologically plausible functional connectivity patterns, relative to pre-existing acquisitions. The protocol and processing pipeline described here are openly shared with the neuroimaging community to promote interoperability and cooperation towards tackling the most important challenges in neuroscience.

## Introduction

Understanding the brain requires a multilevel approach across spatial and temporal scales. Distinct brain network features, as revealed by task-free functional magnetic resonance imaging (fMRI), play a central role in our comprehension of healthy brain function and disorder mechanisms. Human neuroimaging has made great strides in our understanding of the brain through data-sharing initiatives^1–5^. Nonetheless, animal models, particularly small rodents, continue to play an important role in neuroscience discovery, in part, due to the feasibility of performing invasive and terminal manipulations on genetically controlled animals^6^. Specifically, rats are commonly used in pharmacological studies owing to similarities in drug metabolism, as well as in behavioral neuroscience due to their higher proficiency in learning more complex tasks.

Human neuroimaging sharing initiatives have led to a standardization of fMRI acquisition protocols that aid in the dissemination, aggregation and reuse of data^4,7,8^. In contrast, preclinical neuroimaging essentially remains without harmonizing guidelines^9^. Acquisitions in animals are performed under diverse protocols that span different strains, restraint and anesthesia conditions, radiofrequency coil designs, and magnetic field strengths. These impact the generalization of the results and conclusions. Efforts to propose acquisition and / or preprocessing protocols rarely extend beyond the confines of single laboratories, thus limiting interoperability and widespread adoption^9,10^. Thanks to the potential of fMRI in rodents to study the biological basis for connectivity phenomena across the whole-brain longitudinally^11^, an optimized consensus protocol could potentiate future scientific discoveries.

In this preregistered study, we set out to aggregate and make publicly available representative datasets with various fMRI acquisition protocols in the rat and identify experimental parameters associated with robust and reliable functional connectivity detection. We curated the *Multirat_rest* collection (646 rats from 65 datasets) representing protocols used at 46 institutions. Based on the outcome of the analysis of the *Multirat_rest* collection, we devised a new consensus protocol and used it to aggregate the *StandardRat* collection (209 rats from 21 datasets). Preprocessing and confound correction was tailored to rodent data of different characteristics using a rodent-adapted fMRI preprocessing and analysis tool. Our primary outcome was the detection of plausible functional connectivity patterns corresponding to the biologically expected models. Collating data from 50 centers and 855 rats, we show that standardized acquisition and the associated preprocessing pipeline optimizes the detection of distributed fMRI networks in rats. In line with large-scale studies from other species^12,13^ we have freely released all data and the generated code.

## Methods

### Preregistration, code, and data availability

The study was pre-registered (https://doi.org/10.17605/OSF.IO/EMQ4B). Jupyter notebooks demonstrating the analysis code are available under the terms of the Apache-2.0 license (https://github.com/grandjeanlab/MultiRat). The raw datasets are available here: Unstandardized resting-state fMRI (*MultiRat_rest*) (https://doi.org/10.18112/openneuro.ds004114.v1.0.0); Standardized resting-state fMRI (*StandardRat*) (https://doi.org/10.18112/openneuro.ds004116.v1.0.0). The preprocessed volumes, time-series, and quality control files are available here (https://doi.org/10.34973/1gp6-gg97). Image preprocessing, confound correction and connectivity analysis were performed using RABIES 0.3.5 (https://github.com/CoBrALab/RABIES).

### Preregistration deviations

We used the SIGMA rat template^14^ instead of the Papp et al. 2014 template^15^, due to fewer artifacts, additional relevant assets, and improved *in vivo* contrast. Some datasets had field-of-view cropped to ease image registration. Some datasets had time-series cropped to ease the computational load. We used temporal signal-to-noise instead of signal-to-noise ratio as these were shown to be correlated^12^. Detailed deviations are listed here: https://github.com/grandjeanlab/MultiRat.

### Animals

All acquisitions were performed following approval from the respective local and national authorities. Participating laboratories were instructed to provide n = 10 rat imaging acquisitions consisting of one anatomical and one resting-state functional run. Exclusion criteria were unsuitability for RABIES preprocessing (*e*.*g*., dedicated image reconstruction needs, restricted field-of-view). The *MultiRat_rest* collection consists of N = 65 datasets from 46 research centers, n = 646 rats (141/505 f/m). *StandardRat* consists of N = 21 datasets, n = 209 rats (93/116 f/m) from 20 research centers.

### Standardized fMRI acquisition protocol

The standardized protocol was determined based on the outcomes of the analysis of *MultiRat_rest* and used to acquire the *StandardRat* dataset. Acquisitions were performed chiefly in ∼2 months old free-breathing Wistar rats, mixed-sex, and anesthetised using 4% isoflurane and 0.05 mg/kg medetomidine s.c. bolus for induction, and 0.4% isoflurane and 0.1 mg/kg/h medetomidine s.c. for maintenance. Imaging with a gradient-echo echo-planar imaging technique was conducted 40 min post-anesthesia induction, with repetition time = 1000 ms, echo time / flip angle / bandwidth defined as a function of field strength (**Table S1**), repetitions = 1000, matrix size [64 × 64], field-of-view (25.6 × 25.6) mm^2^, 18 interleaved axial slices of 1 mm with 0.1 mm gap. The full protocol is available here: https://github.com/grandjeanlab/StandardRat.

### Data preprocessing and confound correction

Scans were organized according to the BIDS format^16^. Preprocessing was performed on each scan session separately using a reproducible containerized software environment for RABIES 0.3.5 (Singularity 3.7.3-1.el7, Sylabs, California, USA). The preprocessing was performed using autobox^17^, N4 inhomogeneity correction^18^, motion correction^18^, a rigid registration between functional and anatomical scans^18^, non-linear registration between anatomical scan and template, and a common space resampling to (0.3 × 0.3 × 0.3) mm^3^. Visual inspection was performed on preprocessing outputs for all scans for quality control. Five confound correction models were tested, using three approaches based on ICA-AROMA^19^, white-matter and ventricle signal, or global signal regression (**Table S2**). These were done together with motion regression, spatial smoothing to (0.5 mm)^3^, a high-pass filter of 0.01 Hz, and a low-pass filter of either 0.1 or 0.2 Hz.

### Data analysis

To determine functional connectivity in individual rats, seed-based analysis was performed with RABIES in template-space using spherical seeds of 0.9 mm diameter located on the S1 barrel field area (S1bf) and anterior cingulate area (ACA). Functional connectivity was calculated as the Pearson’s correlation coefficient between regional time-courses. Functional connectivity specificity was defined relative to the left S1bf seed, using the contralateral right S1bf region-of-interest as the specific region-of-interest, and the ACA as the unspecific region-of-interest^12^. Functional connectivity was evaluated for each animal and divided into four categories: specific (r_S1bf left to right_ > 0.1 AND r_S1bf left to ACA_ < 0.1), unspecific (r_S1bf left to right_ > 0.1 AND r_S1bf left to ACA_ > 0.1), no (r_S1bf left to right_ [-0.1, 0.1] AND r_S1bf left to ACA_ [-0.1, 0.1]), and spurious connectivity (remaining cases).

### Statistical analysis

One sample t-test voxel-wise maps and group independent component analysis were estimated using Nilearn 0.7.1^20^. Comparisons between functional connectivity specificity and categorical variables (e.g., magnetic field strength, strain, sex) were determined using χ^2^ tests, as implemented in SciPy 1.6.2^21^. Continuous variables (e.g., mean framewise displacement) were transformed into six categorical bins to allow comparison with χ^2^ tests. Linear regression and ANOVA were performed using Pingouin 0.5^22^. Individual seed-based maps are represented as color-coded overlays thresholded at r > 0.1. Given the emphasis on detection of functional connectivity, we mitigated against false negatives by applying a threshold of p_uncorrected_ < 0.05 to the one-sample t-test maps, following preregistration specifications. Slice positions are indicated in mm relative to the anterior commissure.

## Results

We aggregated the *MultiRat_rest* collection of unstandardized fMRI datasets representative of local site acquisition procedures (N = 65 datasets, n = 646 rats). As expected, we found high heterogeneity in all experimental factors recorded, including rat characteristics (sex, strain, **Figure 1a,b**, age, weight, **Figure S1**), in-scan physiology (anesthesia/awake, breathing rates **Figure 1c,e**), and image acquisitions (magnetic field strength, sequence, and sequence parameters **Figure 1d,f,g**). Notably, there was a large sex bias in favor of males (**Figure 1a)**. Despite the heterogeneous distribution of the acquisition parameters and ensuing image quality (**Figure 1g,h,i**), 638/646 of the scans passed preprocessing quality control (one scan with excessive motion, one empty scan, six scans failed image registrations, **Figure S2**). As further quality controls, we described temporal signal-to-noise (**Figure S3**) and motion parameters (**Figure S4**). Overall, we found that the aggregated datasets represent current rodent fMRI acquisition trends^9^ and that the RABIES toolbox can be effectively employed for the preprocessing of rat datasets despite widely varying acquisition parameters. This paves the way for reproducible and interoperable data processing across sites.

**Figure 1.**
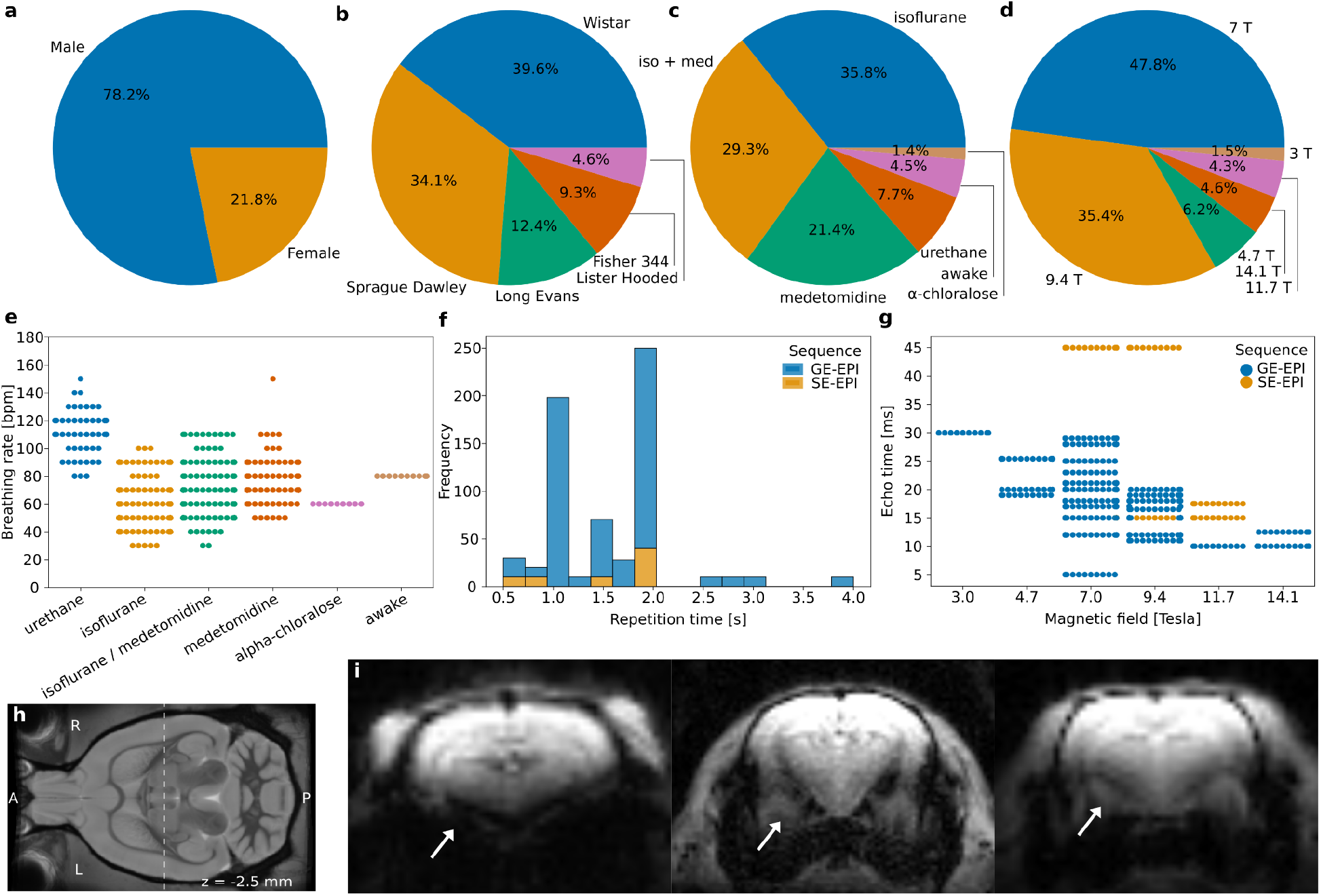
MultiRat_rest dataset description. **a**. Sex, **b**. Strain, **c**. Anesthesia, **d**. Magnetic field strength, **e**. Breathing rate as a function of anesthesia, **f**. Repetition time, **g**. Echo time as a function of magnetic field strength, **h**. Slice position for the examples, **i**. Example of representative raw functional images. Arrows indicate different susceptibility artifact-related geometric distortions in the amygdala.

We focused on examining functional connectivity in the sensory cortex, as sensory networks are robust to anesthesia effects, in the anesthesia depth range typically used in fMRI^23^. More specifically, we evaluated the specificity of the connectivity of the S1 barrel field area (S1bf) using two complementary criteria as indexes of accurate functional connectivity identification (**Figure 2a,b**)^12^. The first criterion was the strong connectivity between inter-hemispheric sensory cortices (barrel field, S1bf). Indeed, in both humans and animals, dating back to the original description of functional connectivity^24^, the majority of the networks including sensory-motor networks have a bilateral homotopic organization. The second criterion was a weak or anti-correlation between S1bf and the anterior cingulate area (ACA). The ACA is a major node in the task-negative rodent default-mode network. Task-positive (as the S1bf-associated sensory network^25,26^), and task-negative networks are generally non- or anti-correlated^27^.

**Figure 2.**
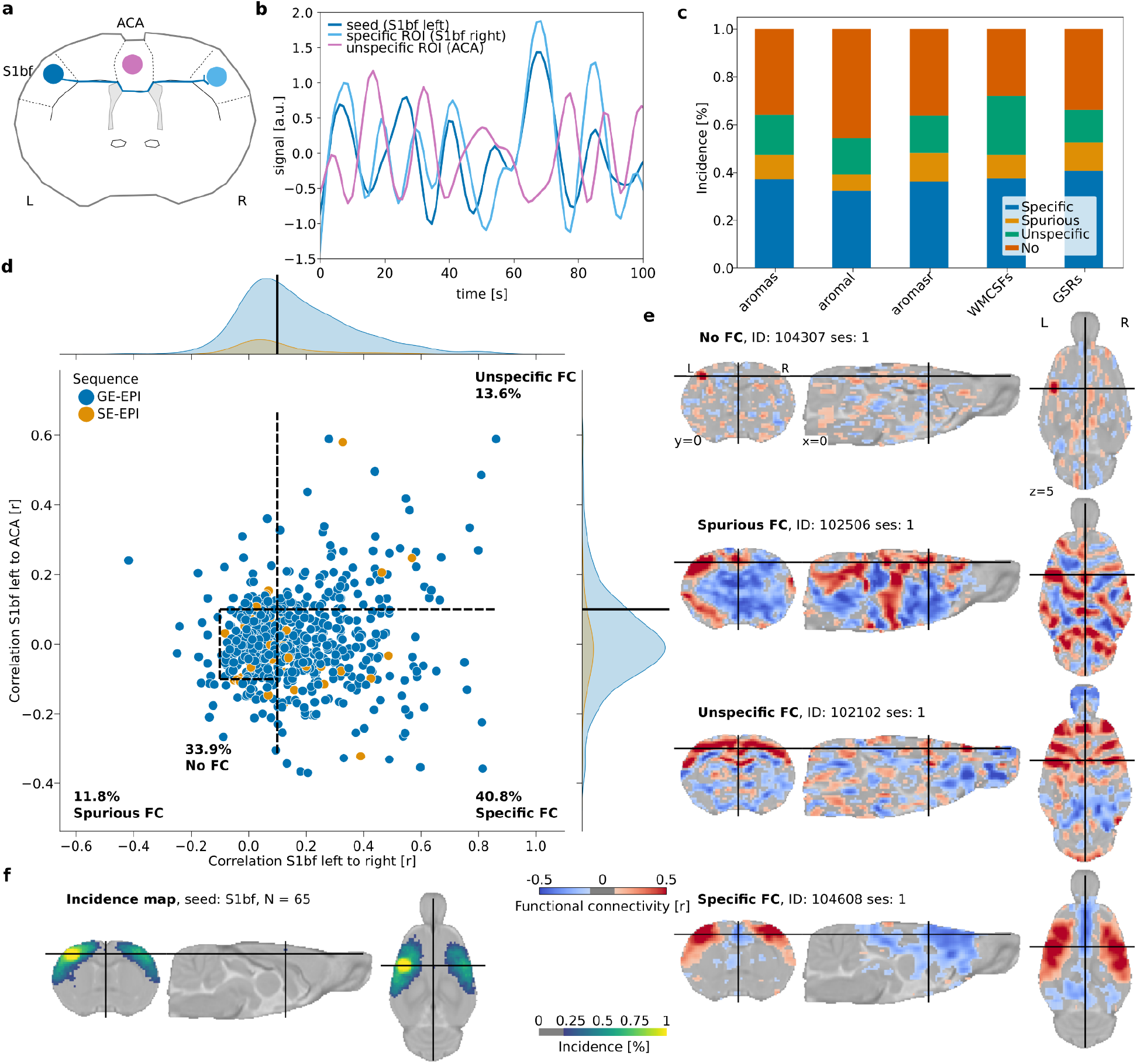
Functional connectivity specificity. **a**. Diagram illustrating the logic behind functional connectivity specificity. The sensory (barrel field, S1bf) area (blue) chiefly projects to the contralateral homotopic area (light blue), but not to the anterior cingulate area (ACA) area (purple) **b**. Example of temporal dynamics in the resting-state signal. Correlated signal between the ipsi- and contralateral S1bf, and anti-correlated signal from the ACA. **c**. Distribution of functional connectivity (FC) categories as a function of confound correction models. **d**. Functional connectivity in left S1bf relative to specific (right S1bf) and unspecific (ACA) ROIs using the global regression correction model. Dots represent scans (n = 638 rats), dotted lines indicate the thresholds used to delineate the categories. **e**. Example of individual seed-based analysis maps for each connectivity category. **f**. Group-level functional connectivity incidence map (N = 65 datasets).

Functional connectivity was evaluated for each animal and divided into four categories based on these two criteria referred above: specific, unspecific, spurious, and no connectivity. Five confound correction models were tested (**Table S2, Figure 2c**). The global signal regression nuisance model was the one that performed the best for specific connectivity detection (40.8% of the animals with specific connectivity, 11.8% as unspecific, 13.6% as spurious, and 33.9% as containing no detectable functional connectivity, **Figure 2c,d,e**). Because network inference is often assessed at the group-level rather than at the individual-level, we performed a one-sample t-test per dataset to estimate the incidence of contralateral connectivity detected within groups, relative to the S1bf seed (**Figure 2f**). Up to 70% of the 65 datasets presented limited evidence of contralateral connectivity relative to the seed, and 50% of the datasets captured the features of a larger sensory network at the group level (see also other seeds, **Figure S5**). We conclude that rat datasets do not capture functional connectivity equally, similar to what we previously reported in the mouse^12^. Although we found that global signal regression enhances the incidence of specific connectivity of S1bf seeds, we can not generalize this nuisance model to examine other connectivity outcomes. Indeed, the global signal partakes in previously unsuspected roles in the signal acquired in task-free paradigms^28^, and its removal should be considered with care.

Our observations underline the need for an improved acquisition protocol to maximize individual-level inferences to potentiate discovery in experimental network neuroscience. To address this, we evaluated parameters associated with increased specificity incidence in the *MultiRat_rest* collection. Medetomidine/isoflurane anesthesia combination condition was enriched in scans categorized as specific (**Figure S6a**, 92/187 scans, χ^2^ test: **φ** = 0.27, dof = 15, g = 92.38, p = 3.5e-13). The use of gradient echo imaging sequence was also associated with higher specificity incidence (**Figure S6b**, 241/568 scans, χ^2^ test: **φ** = 0.11, dof = 3, g = 16.00, p = 0.001). Based on these observations, we devised an anesthesia protocol derived from dataset ds01031 (9/10 specific scans), and an imaging sequence based on ds01028 (8/10 specific scans), acquired on a 4.7 T mid-field system. We hypothesized that this protocol would enhance functional specificity while being compatible with lower field systems that continue to represent a relevant share of the systems in use (**Figure 1d**)^9^.

Using this consensus protocol, we curated the *StandardRat* collection of 21 datasets obtained across 20 centers. This consisted of n = 209 rats (93/116 f/m) rats, chiefly Wistar (189/209) aged ∼ 2 months (**Figure S7**). Dataset acquisitions were performed at magnetic field strengths ranging from 4.7 to 17.2 T. Preprocessing was performed similarly to the unstandardized dataset. 207/209 scans passed quality assurance (two discards due to image misregistration). Interestingly, despite the same anesthesia protocol being used, the respiratory rates reported at the start of fMRI acquisition differed as a function of rat strain (**Figure 3a,b**, ANOVA, η^2^ = 0.24, F_(195,2)_ = 31.17, p = 1.8e-12,). Finally, there was only a negligible effect on the temporal signal-to-noise ratio as a function of magnetic field strength (**Figure 3c**, linear regression, coef = 0.53 [-0.23, 1.30], r^2^ = 0.01, dof = 201, T = 1.37, p = 0.17).

**Figure 3.**
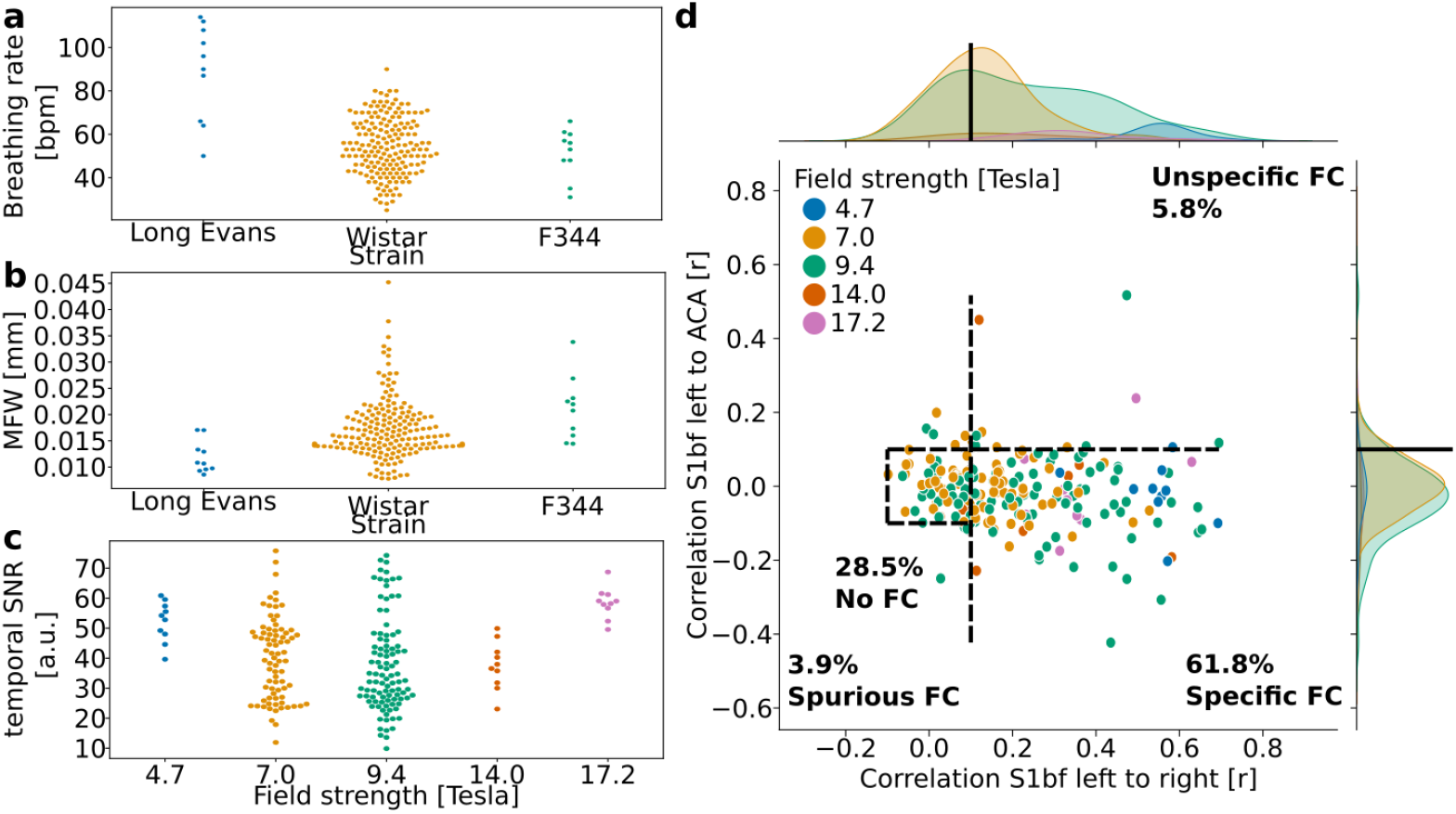
StandardRat dataset description. **a**. Breathing rate (breath per minute, bpm) as a function of strain. **b**. Mean framewise displacement (MFW) as a function of strain. **c**. Temporal signal-to-noise ratio in the sensory cortex as a function of field strength. **d**. Functional connectivity in left S1bf relative to specific (right S1bf) and unspecific (ACA) ROIs using the global regression correction model. Dots represent scans (n = 207 rats), dotted lines indicate the thresholds used to delineate the categories.

The objective for this study was to find an improvement in specific connectivity in the individual datasets. We found that 61.8% of the scans were categorized to contain specific connectivity (**Figure 3d)** against 40.8% in the *MultiRat_rest* dataset with unstandardized acquisitions (**Figure 2d)** when using global signal regression (χ^2^ test: **φ** = 0.13, dof = 3, g = 33.01, p = 3.2e-07). The difference remained when we compared datasets from centers that contributed to both collections exclusively (χ^2^ test: φ = 0.17, dof = 3, g = 28.37, p = 3.0e-06). This is going against the notion that the *StandardRat* collection is outperforming due to an enrichment in datasets from more experienced laboratories. Intriguingly, we could not establish a field strength effect on connectivity specificity (χ^2^ test: **φ** = 0.19, dof = 12, g = 14.89, p-value = 0.25), suggesting acquisition systems are not the limiting factor in this protocol. We conclude that the newly standardized protocol outperforms, on average, previously used protocols within the community for the detection of biologically plausible connectivity patterns.

Intriguingly, there remained differences in the connectivity patterns between datasets from the *StandardRat* collection. We next sought to identify the variables associated with greater incidences of specific connectivity patterns. Importantly, we could not establish strain, sex, or magnetic field strength effects, suggesting the protocol is applicable for a large range of conditions (**Table S3**). Next, we examined breathing rate and temporal signal-to-noise ratio as indicators of acquisition variability (**Figure S8**). Overall, we found that scans with breathing rates ranging 84-114 breaths-per-minutes, cortical temporal signal-to-noise ratio >53 achieved higher incidences of connectivity specificity among the *StandardRat* collection. These provide the first line of evidence to refine the *StandardRat* protocol by identifying practices that can further enhance connectivity outcomes.

In summary, we curated two dataset collections (*Multirat_rest* and *StandardRat)*, analyzed them and made them an open-access resource. These are the largest rodent fMRI datasets currently available. We developed and deployed a preprocessing and confound correction strategy generalizable to most scans and every dataset. Using information from the *MultiRat_rest* collection, we provide useful population parameter estimates to enhance the comparison of rat fMRI datasets. We proposed and evaluated a new standardized protocol and found that this consensus acquisition and preprocessing pipeline outperformed the previous acquisitions for connectivity specificity. To allow replication and to inspire new analyses we release all raw and processed data to the broader community.

On average, the standardized protocol yielded improvements over previous acquisitions gathered in the *MultiRat_rest* collection. However, individual-level inferences remain limited to 61.8% of the scans acquired. This underlines the importance of implementing sound quality assurance metrics based on assumptions of biologically-plausible functional connectivity. Improving output quality, either through understanding the factors leading to successful acquisitions, enhanced protocols, preprocessing, or nuisance regression models would lead to tangible outcomes capable of further potentiating future data acquisitions and reducing animal use by reducing discards.

Importantly, our new protocol relies on light sedation to restrain the animals. While optimized for fMRI, this protocol may not generalize to other procedures such as electrophysiology. We also found that existing awake restraining protocols, on average, lead to lower incidence of specific connectivity patterns. A previous report has indicated similar values in a dataset in awake rats^29^. Due to the impact of anesthesia on networks^9,10^, it remains central to develop awake imaging as an alternative. However, these protocols should be examined through the lens of quality control metrics to ensure plausible connectivity patterns are achieved consistently. Further, the acquisition sequence in *StandardRat* is designed to run on a wide range of systems. The effectiveness of new sequences should be examined against the current protocol, e.g., isotropic resolution^30^ or multiband acquisition^31^.

Our project’s methodological and conceptual advancements are the first step towards large multi-site rat neuroimaging acquisitions. Coordinated open-science projects in neuroimaging and other disciplines are transforming the scientific landscape^32^. Through the concerted efforts of our centers and potentiated by a substantially improved protocol, rat functional brain imaging is set to tackle urgent questions in neuroscience and mental health research.

## Acknowledgments

This research was enabled in part by support provided by Compute Ontario https://www.computeontario.ca/ and Compute Canada (www.computecanada.ca). For the purpose of Open Access, the author has applied a CC BY public copyright licence to any Author Accepted Manuscript version arising from this submission.

