## Supplementary material for "StandardRat: A multi-center consensus protocol to enhance functional connectivity specificity in the rat brain"

**Table S1 | Magnetic field strength-dependent parameters for the standardRat acquisition protocol.**

| | **3T** | **4.7T** | **7T** | **9.4T** | **11.7T** | **14.1T** | **17.2T** |
|---|---|---|---|---|---|---|---|
| **TE [ms]** | 30 | 25 | 17 | 15 | 12 | 10 | 9 |
| **Flip angle [degree]** | 64 | 61 | 55 | 53 | 52 | 51 | 50 |
| **Receiver bandwidth [kHz]** | 180 | 200 | 220 | 250 | 250 | 300 | 300 |

**Table S2 | Confound correction model.**

| **Confound model** | **Low-pass filter** | **Nuisance regression** | **Other** | **RABIES arguments** |
|---|---|---|---|---|
| *aromas* | 0.1 Hz | ICA-AROMA | - | --lowpass 0.1 --run_aroma --aroma_dim 10 |
| *aromal* | 0.2 Hz | ICA-AROMA | - | --lowpass 0.2 --run_aroma --aroma_dim 10 |
| *aromasr* | 0.1 Hz | ICA-AROMA | Framewise displacement censoring together with 1 back and 2 forward frames at FD threshold 0.05 | --lowpass 0.1 --run_aroma --aroma_dim 10 --FD_censoring |
| *WMCSFs* | 0.1 Hz | White matter + Cerebrospinal fluid + motion | - | -lowpass 0.1 --conf_list WM_signal CSF_signal mot_6 |
| *GSRs* | 0.1 Hz | Global signal regression + | - | --lowpass 0.1 --conf_list |

| | | | | |
|---|---|---|---|---|
| | | motion | | global_signal<br>mot_6 |

**Table S3 |** Frequency test ($\chi^2$) for connectivity categories in the StandardRat collection

| Effect | Effect size (**φ** ) | Degrees of freedom | g-value | p-value |
|---|---|---|---|---|
| **Strain** | 0.15 | 6 | 8.77 | 0.19 |
| **Sex** | 0.07 | 3 | 2.16 | 0.54 |
| **Field strength** | 0.19 | 12 | 14.89 | 0.25 |

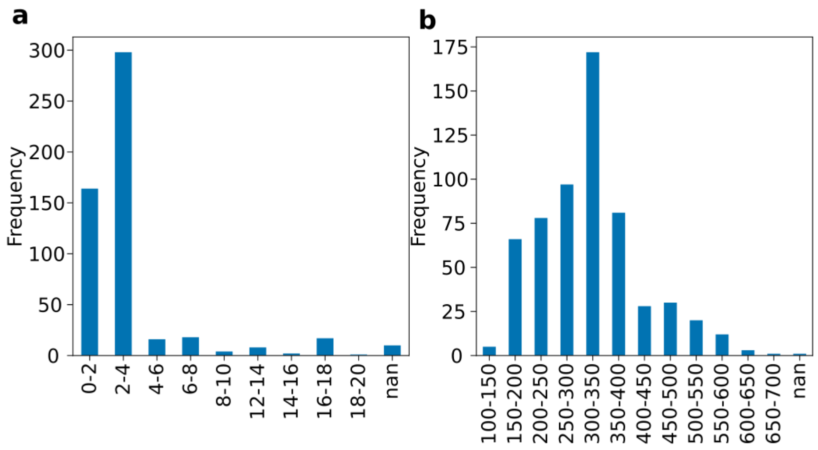


**Figure S1**. *Age (**a**) and weight (**b**) distribution for the rats in the multiRat_rest collection.*

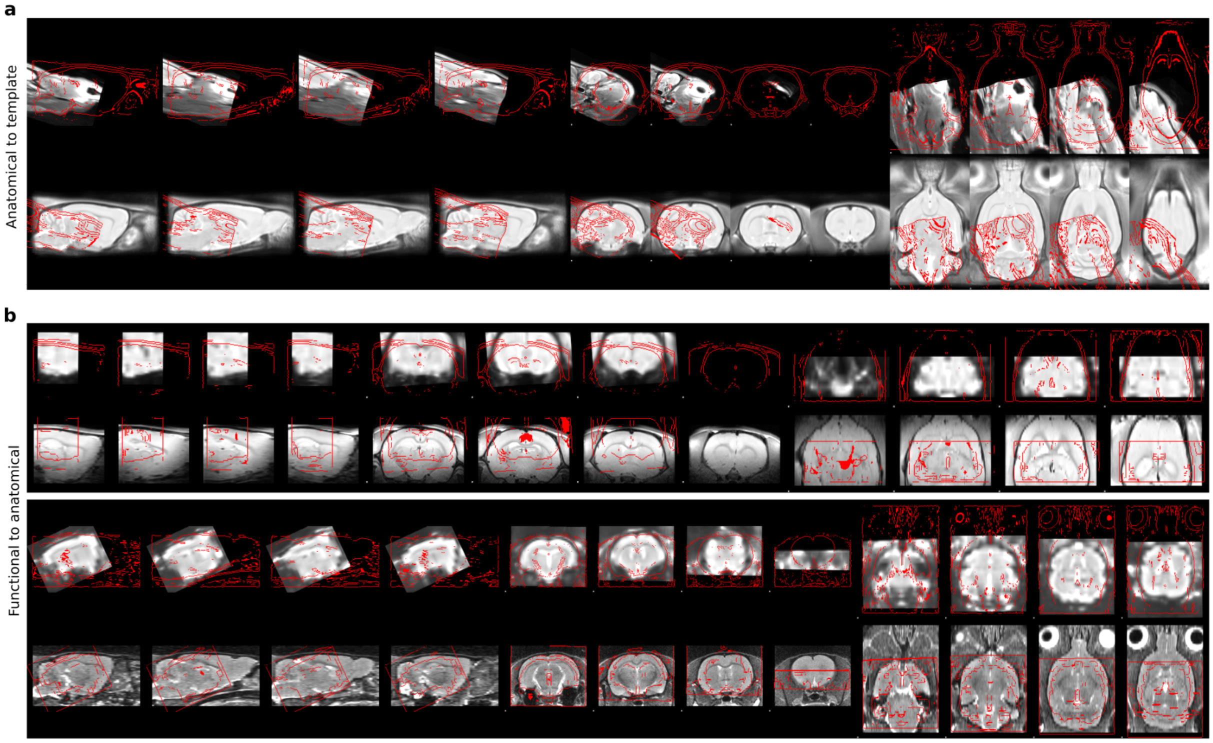


**Figure S2**. *Failed quality controls for anatomical to template registrations (**a**) and functional to anatomical registrations (**b**). The top rows are the moving objects, bottom rows are the reference objects. Vice versa for the red outlines.*

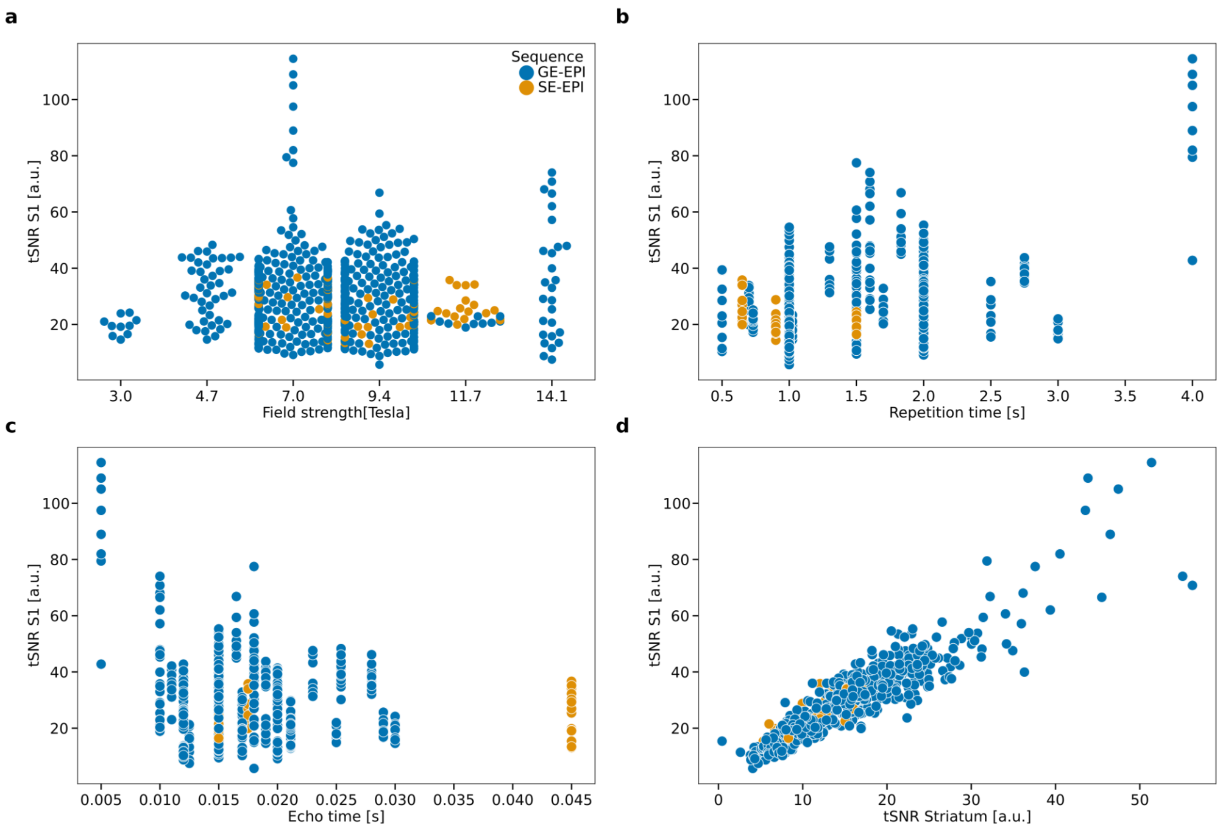


**Figure S3**. *Temporal signal-to-noise ratio in the sensory cortex (tSNR S1) in the multiRat_rest dataset collection as a function of (**a**) magnetic field strength, (**b**) repetition time, (**c**) echo time, (**d**) temporal signal-to-noise ratio in the striatum.*

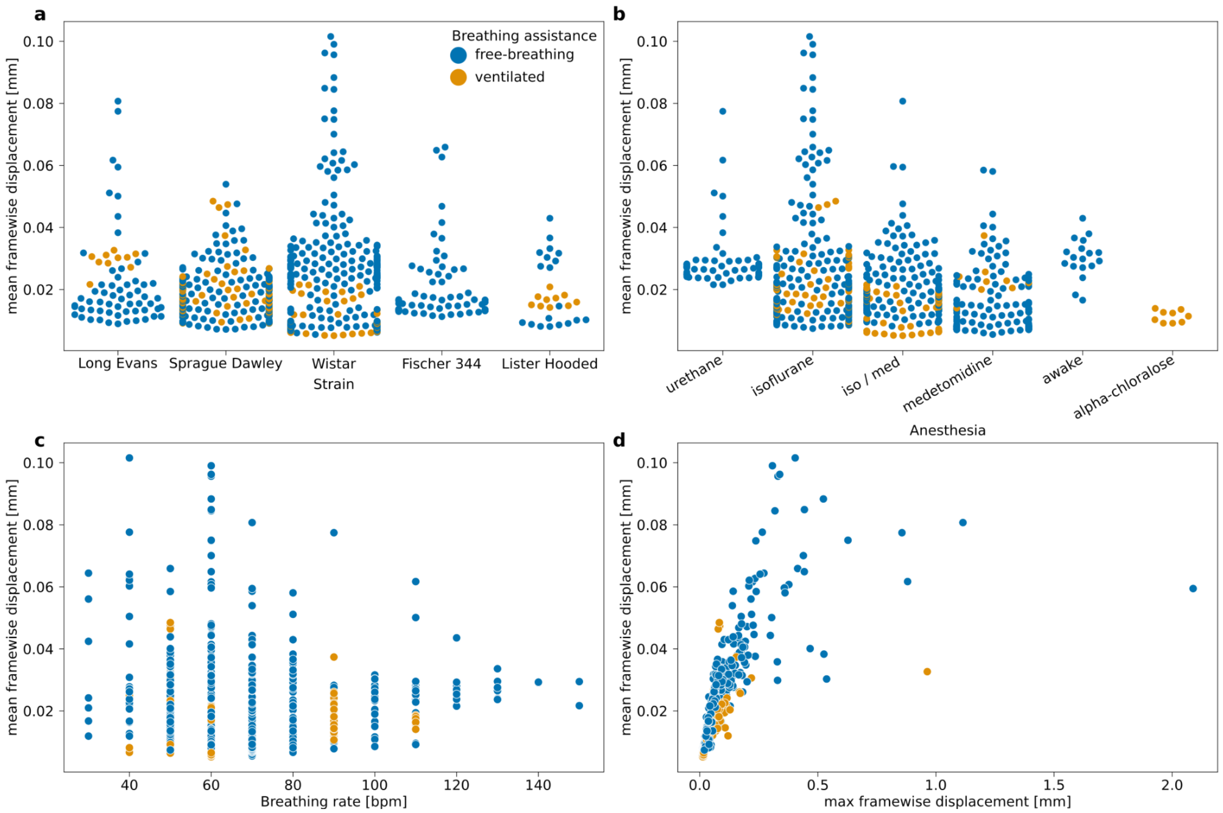


**Figure S4**. *Mean framewise displacement in the multiRat_rest dataset collection as a function of (**a**) strain, (**b**) anesthesia, (**c**) breathing rate, (**d**) maximal framewise displacement.*

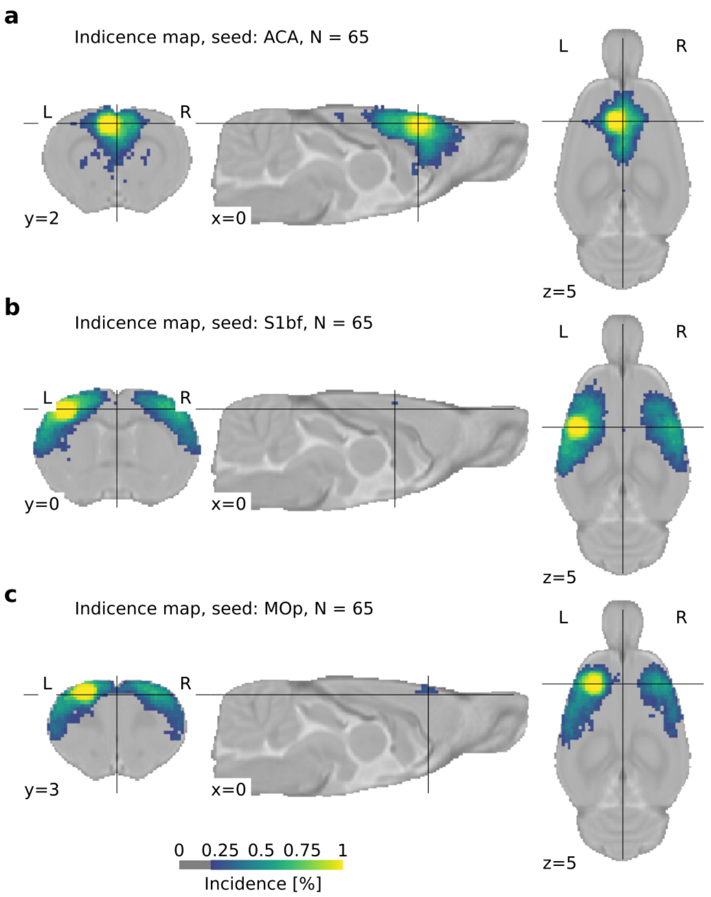


**Figure S5**. *Incidence of functional connectivity at the group level (N=65 datasets of n ~ 10 subjects per dataset). Similar to functional connectivity relative to the sensory barrel field seed (***Figure 2f***), distal connectivity relative to the seed, either along the AP-axis for the anterior cingulate area (ACA) seed, or contralateral for the primary motor area (MOp) and caudoputamen (CPu) seeds, were generally observed in 50 - 75% of the datasets.*

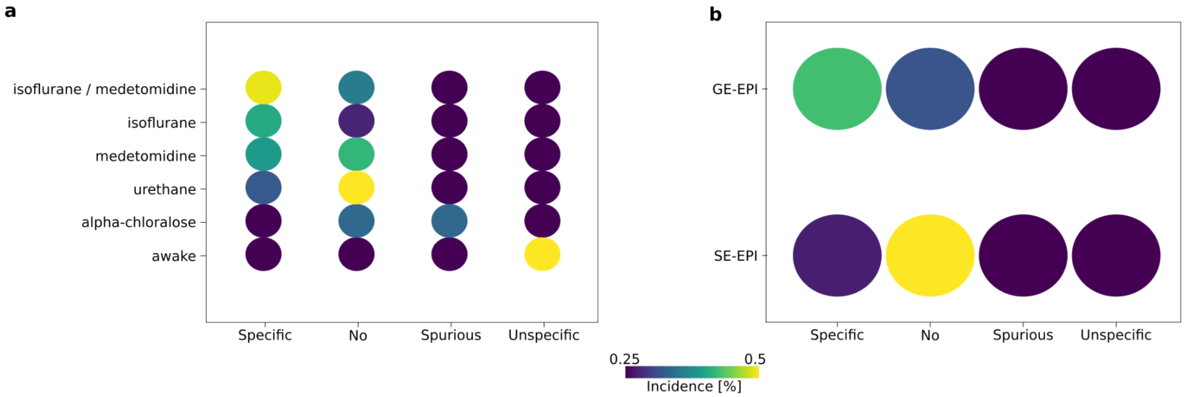

**Figure S6**, *Functional connectivity specificity as a function of anesthesia (**a**) and acquisition sequence (**b**) in the mutliRat_rest collection. A greater percentage of scans achieve connectivity specificity when the anesthesia is maintained with isoflurane combined with medetomidine. More scans achieved the same endpoint when acquired with gradient-echo echo planar imaging (GE-EPI) than spin-echo (SE-EPI). The percentage of each condition is color-coded.*

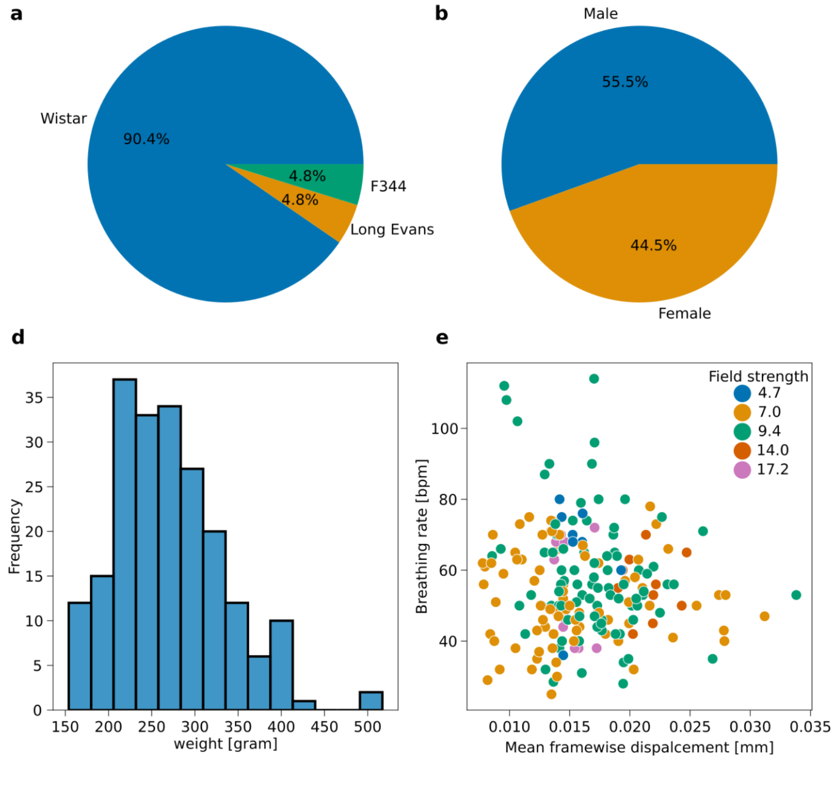


**Figure S7**. *StandardRat dataset description.* ***a****. Strain.* ***b****. Sex.* ***c****. Field strength.* ***d.*** *Weight.* ***e.*** *Breathing rate as a function of mean framewise displacement.* ***f.*** *Functional connectivity specificity as a function of confound correction models.*

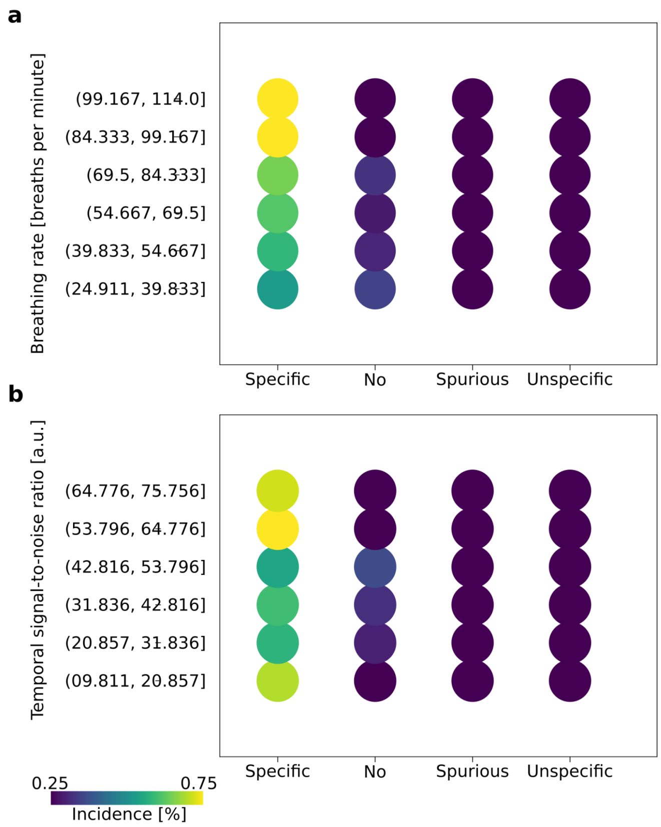


**Figure S8**, *Functional connectivity specificity as a function of binned breathing rate (**a**) AND temporal signal-to-noise ratio (**b**) in the standardRat collection.The percentage of each condition is size and color-coded. High levels of connectivity specificity were achieved in scans where the breathing rates were in the 84 to 114 bpm range. Similarly, higher connectivity specificity incidences were found when the cortical temporal signal-to-noise ratio was >53. These observations support the notion of an optimal breathing rate when applying the standardRat protocol, along with temporal signal-to-noise ratio and movement targets.*
